# A quantitative tri-fluorescent yeast two-hybrid system: from flow cytometry to *in-cellula* affinities

**DOI:** 10.1101/553636

**Authors:** David Cluet, Ikram Amri, Blandine Vergier, Jérémie Léault, Clémence Grosjean, Dylan Calabresi, Martin Spichty

## Abstract

We present a technological advancement for the estimation of the affinities of Protein-Protein Interactions (PPIs) in living cells. A novel set of vectors is introduced that enables a quantitative yeast two-hybrid system based on fluorescent fusion proteins. The vectors allow simultaneous quantification of the reaction partners (Bait and Prey) and the reporter at the single-cell level by flow cytometry. We validate the applicability of this system on PPIs with different affinities. After only two hours of reaction, expression of the reporter can easily be detected even for the weakest PPI. Through a simple gating analysis, it is possible to select only cells with identical expression levels of the reaction partners. As a result of this standardization of expression levels, the mean reporter levels directly reflects the affinities of the studied PPIs. With a set of PPIs with known affinities, it is straightforward to construct an affinity ladder that permits rapid classification of PPIs with thus far unknown affinities. Conventional software can be used for this analysis. To permit automated high-throughput analysis, we provide a graphical user interface for the Python-based FlowCytometryTools package.

## Introduction

Protein-protein interactions (PPIs) are essential for many functions in living cells, including communication of signals, modulation of enzyme activity, active transportation, or stabilization of the cell structure by the cytoskeleton (1–4). Resolving the complex cellular network of PPIs remains one of the major challenges in proteomics (5). Thus, the quest for reliable methods that identify PPIs and quantify their strength is unbroken.

The yeast two-hybrid technique (Y2H) is a commonly used approach to probe the interaction between proteins (6–8). In contrast to biochemical *in-vitro* methods (such as mass spectrometry, ITC or SPR) that require purified proteins, Y2H is based on a genetic assay. It relies on the *in-cellula* expression of fusions of the two proteins of interest, usually named Bait and Prey. Upon physical interaction of Bait and Prey, a functional transcription factor is reconstituted that drives the expression of a reporter gene (*e.g.*, β-galactosidase). As a consequence, a read-out is observed (e.g., color, fluorescence, or growth) that permits high-throughput screens.

Y2H has been extensively used in the past decades to decipher PPI networks (9, 10). With growing experience, the scientific community became aware of the limitations of this approach. Standard Y2H is prone to false positive/negative results (8). For example, the absence of a detectable read-out may reflect insufficient expression of the Bait and/or Prey. More laborious Western blottings can be performed to verify the expression (11). Furthermore, Y2H provides often only a qualitative result. With X-gal-based Y2H (12), for instance, the measured read-out (color) cannot be assumed to be proportional to the reporter level, *i.e.*, β-galactosidase activity, but exceptions exist (13, 14). Also, the extent of β-galactosidase activity does not necessarily reflect the extent of interaction between Bait and Prey (due to varying expression levels of Bait and Prey fusions).

Several groups tried to overcome the qualitative limitations of the two-hybrid system in yeast and other organisms. Extensive overviews can be found in the literature, for example Ref. (8). Many applied methods could rank PPIs according to their affinity using the quantified read-out, examples are included in Refs. (13–17). It should be noted, however, that mainly mutants were compared. Similar expression levels for the Bait and Prey fusions can be assumed for such mutational studies. Comparing proteins from different families often breaks the correlation (11). It speaks to the need of quantifying not only the read-out but the Bait/Prey fusions as well. Thus far, only low-throughput methods exist that address this necessity. For example, by measuring the fraction of co-localized fluorescent “Bait” and “Prey” fusions in human cells by high-resolution microscopy (18) it was possible to optimize the affinity of an inhibitor (19). Another approach used a fluorescent antibody to quantify the amount of retained Prey by the Bait associated to the periplasm (20). Following this idea, different yeast surface two-hybrid approaches emerged (17, 21) using antibodies or purified proteins.

Here we present a novel set of Y2H vectors that enable the quantification of the reaction partners (Bait, Prey) and the reporter at the single-cell level without the need of any antibodies or purified proteins (Fig. 1). Three different fluorescent tags serve as sensors to probe the cellular expression levels by flow cytometry. The vectors have been validated on protein-protein interactions with varying affinities (see Table 1) to encompass the sensitivity of the novel quantitative Y2H (qY2H) system.

**Fig. 1.**
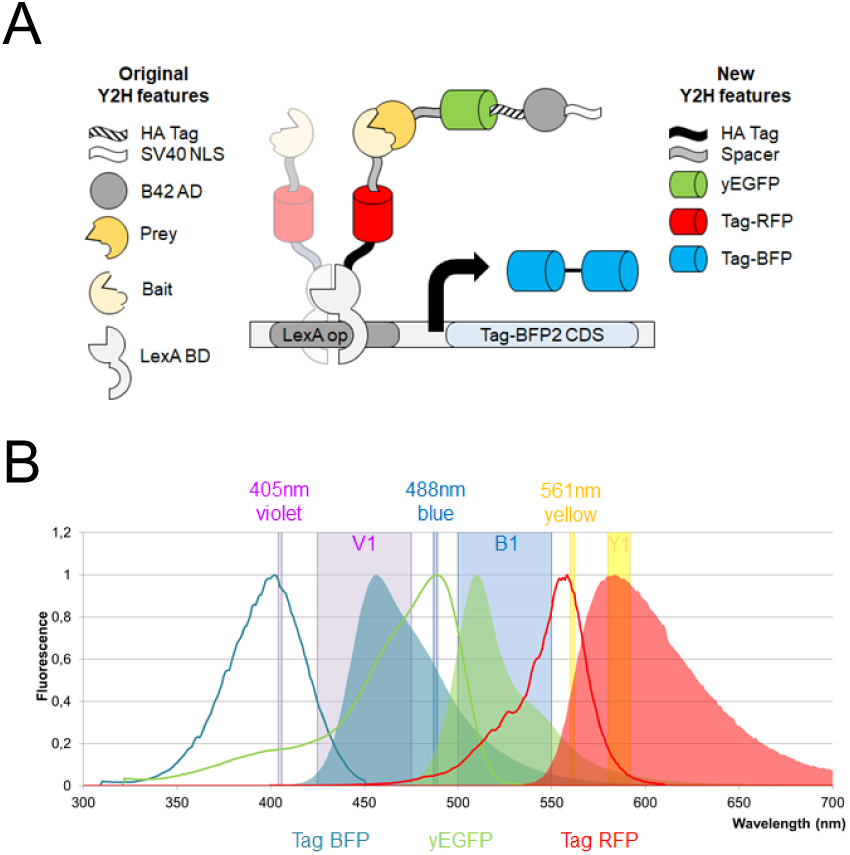
Concept of the qY2H approach. *A.* Our novel Y2H system is based on the one of Brent and coworkers (22). It relies on a split transcription factor where the Bait protein is fused to the LexA DNA-binding domain (BD). The latter binds specifically to the operator of the reporter cassette. The Prey protein is fused to the B42 activation domain (AD). The interaction between BD-Bait and AD-Prey reconstitutes a functional transcription factor that drives the expression of the reporter gene. Our quantitative Y2H approach monitors as a novelty the expression levels of BD-Bait, AD- Prey and the reporter at the single cell level using fluorescent tags. The original fusion proteins when expressed with the constructs of Brent and coworkers are described in the left column of the legend. Our newly added features are presented in the right column. The spacer amino-acid sequence is EFGRALE. *B.* The three fluorescent proteins have separated excitation (bold lines) and emission (areas) spectra. Thus, their expression can be easily monitored using compatible flow cytometers. In this work we used a MacsQuant VYB flow cytometer. The three lasers (Violet, Blue, and Yellow) and their respective detection channels (V1, B1, and Y1) are represented as vertical bands.

**Table 1:**
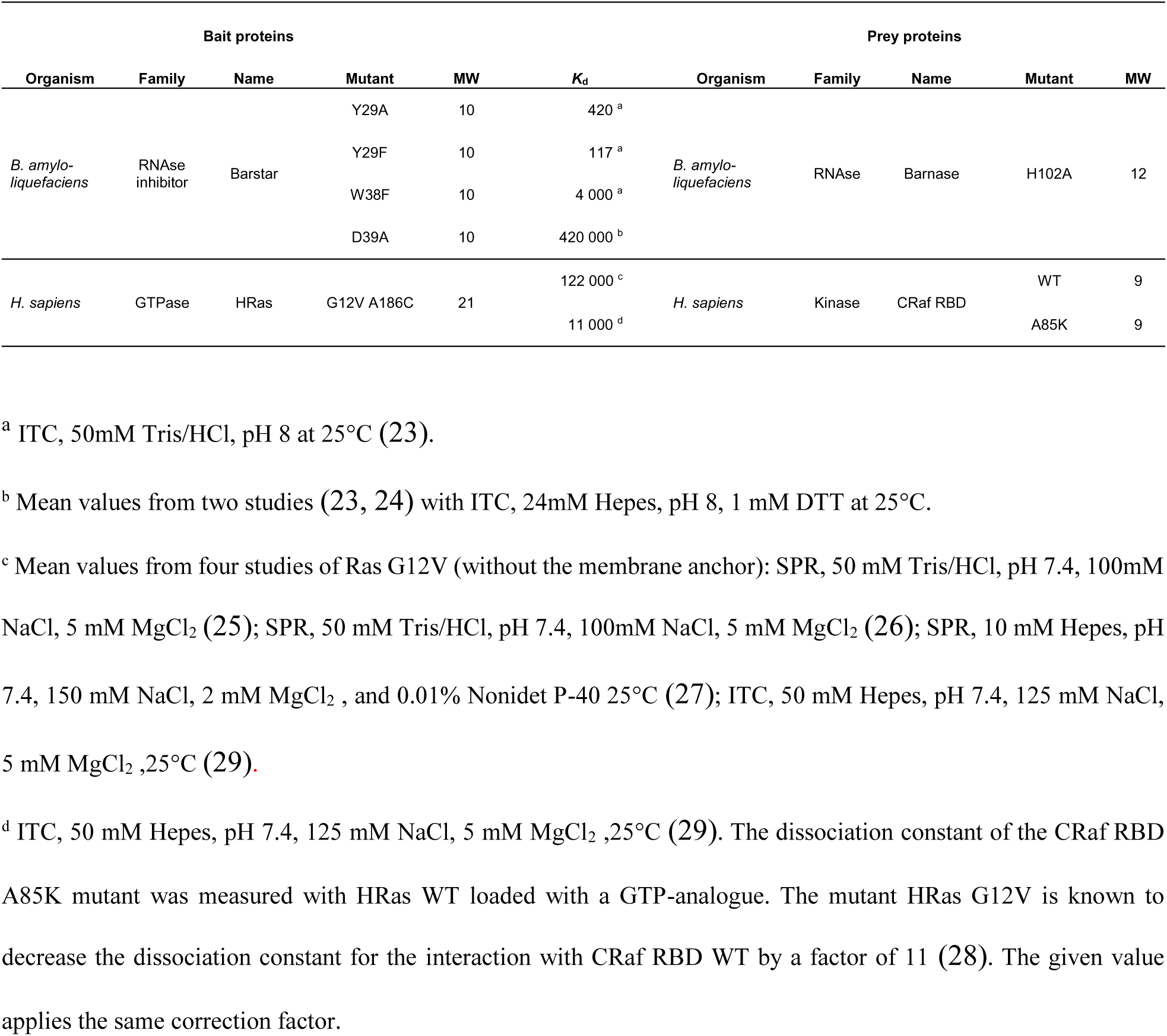
List of BD-Bait and AD-Prey couples tested with the qY2H approach. The dissociation constants (*K*_d_) were taken from the referenced literature and are given in pM; the molecular weights (MW) are in kDa. CENTER DONE

## Experimental Procedures

### Experimental design

Starting from the original plasmids pLexA and pB42AD (22), several new cloning sites were introduced that permit convenient sub-cloning into the expression cassettes via homologous recombination in yeast (see Suppl. Fig. S1 for detailed vector maps). Thus, the newly designed vectors facilitate the construction of novel fusions proteins with tailored functionalities. Here we generated cassettes that code for BD-Bait, AD-Prey and reporter fusions with several new features as shown in Fig. 1. We copied the HA tag (that was originally only in the AD-Prey expression cassette) to the BD-Bait cassette to enable the simultaneous quantification of expressed BD-Bait and AD-Prey fusions by Western Blotting. In addition, we added red and green fluorescent tags (Tag RFP and yEGFP) to the BD-Bait and AD-Prey cassettes, respectively. Furthermore, the original reporter (β-galactosidase) was replaced by a tandem of the Tag BFP in the pSH18-34 vector (22). The three fluorescent tags emit at considerably different wavelength ranges so that their individual expression levels can be simultaneously monitored at the single-cell level by flow cytometry (Fig. 1B). Also, a spacer sequence was added between the fluorescent tags and the Prey/Bait to avoid steric hindrance in the expressed fusions.

The new constructs were tested on two well-studied protein-protein interactions (23–29): the association between the bacterial ribonuclease Barnase and its inhibitor Barstar, as well as between the human GTPase HRas and the Ras-binding domain of CRaf. Different mutants were applied (Table 1) that span a wide range of affinities (known from independent *in-vitro* experiments). By exchanging Bait and Prey for a given couple it is possible to probe the corresponding PPI in two different orientations. We cross-tested all proteins of Table 1 as BD-Bait and AD-Prey fusions with proper negative controls, *i.e.*, empty AD-EGFP and BD-RFP fusions, respectively, to eliminate orientations that lead to false positive interactions (see Material & Methods).

For a given couple, the two orientations may also lead to different read-outs (15). Table 1 lists the couples in the orientation with the stronger reporter level (when the cellular contents of BD-Bait and AD-Prey are standardized, see below). This orientation is considered as the molecular configuration with the higher accessibility of the PPI binding interface (15). Or in other words, this orientation may feature the smaller steric hindrance due to the fused fluorescent BD and AD and probably resembles more closely the situation of free (not fused) proteins. We focus therefore in the following sections on the orientation given in Table 1.

Haploid cells were transformed with either Prey or Bait plasmids; in the latter case we used haploid cells that were previously transformed with the reporter plasmid. Transformed yeast cells were mated and amplified to generate diploid colonies for the desired Bait/Prey-couples. Selection and amplification of the diploids occurred in glucose medium which represses the expression of the AD-Prey fusion (under the control of the GAL1 promoter). Transfer of the diploid cells into Galactose/Raffinose medium induced the expression of the AD-Prey fusion and enabled the expression of the reporter.

After fixation, the samples were submitted to flow cytometry measurements to monitor the fluorescence intensities at the single cell level for three different channels matching the emission ranges of the fluorescent BD-Bait, AD-Prey and reporter fusions; hereafter these channels are named Tag-RFP-H, yEGFP-H and Tag-BFP-H, respectively.

We analyzed the expression level of the fluorescent proteins either for the entire cell population (to which refer as “global” hereafter) or for subpopulations using interval gatings. The entire procedure starting from the transformation up to the flow cytometry measurement was repeated at least three times for each protein-protein interaction.

### Plasmid creation

In order to generate the pSB_1Bait plasmid, the pLexA (22) vector was linearized using *Eco*RI and *Sal*I (Thermo Scientific) to remove all DNA between the LexA cDNA and the ADH terminator. The Barstar WT coding sequence was ordered for synthesis to Eurofin Genomics as part of a new expression cassette. At the 5’ end we added the sequences of the HA-Tag and our MCS-spacer (*Eco*RI, *Asc*I, and *Xho*I). At the 3’ end, after the stop codon of Barstar, we inserted one *Xho*I site, created 3 stop codons (1 per ORF) and regenerated the *Sa*lI site. The upstream (LexA) and downstream (Terminator) 30bp required for homologous recombination in yeasts (30, 31) were also added. This new optimized expression cassette was amplified by PCR (Phusion DNA polymerase, Thermo Scientific), using the primers primSB_0001 and 2 (see Suppl. Table S1), and inserted in the previously linearized pLexA vector. As a result, we obtained the pSB_1Bait_Barstar. The coding sequence for Tag-RFP was subsequently introduced in the *Eco*RI site through PCR from pTag_RFP-Actin (Evrogen), using the primers primSB_0003 and 0004, combined with homologous recombination in yeasts to obtain the pSB_1Bait_RFP-Barstar plasmid. The pSB_1Bait_RFP-Empty and pSB_1Bait-Empty vectors were generated by digesting the pSB_1Bait_RFP-Barstar and pSB_1Bait_Barstar, respectively, with *Xho*I (Thermo Scientific), followed by self-ligation. The coding sequences of the mutants of Barstar, and Ras_G12V_C186A were ordered to Eurofin Genomics, amplified (primSB_0018 and 0019) and introduced in the pSB_1Bait_RFP-Empty linearized with *Xho*I by homologous recombination.

To create the pSB_1Prey vector, the pB42AD plasmid (22) was linearized using *Eco*RI and *Xho*I. The sequence coding for the non-toxic Barnase mutant H102A was ordered from Operon MWG. At the 5’ end we inserted the same MCS-spacer sequence as in the pSB_1Bait vector to allow easy transfer from one plasmid to the other. At the 3’ end, we inserted one *Xho*I site, created 3 stop codons and one *Nco*I restriction site. The upstream (HA-Tag) and downstream (Terminator) 30bp required for homologous recombination in yeasts were also introduced. This new expression cassette was then amplified by PCR (primSB_0010 and 0011) and inserted in the pB42AD by homologous recombination in yeast to obtain the pSB_1Prey_Barnase-H102A vector. The coding sequence of the yEGFP was amplified from the pGY-LexA-GFP_KanMX (kindly provided by Dr Gaël Yvert) using the primers primSB_0012 and 0013, and then introduced in the *Eco*RI sit of our MCS as previously to generate the pSB_1Prey_yEGFP-Barnase-H102A vector. The pSB_1Prey-Empty and pB_1Prey_yEGFP-Empty were created by removing the coding sequence of Barnase H102A with *Xho*I and performing a self-ligation. The coding sequences of the other Preys (CRaf RBD WT, and CRaf RBD A85K) were ordered to Eurofin Genomics, amplified by PCR (primSB_0020 and 21) and introduced in the pSB_1Prey_yEGFP-Empty linearized with *Xho*I by homologous recombination.

To create the reporter plasmid, the pSH18-34 (22) was digested using the unique *Sal*I (In the modified Gal1 promoter) and *Rsr*II (downstream to the β-Galactosidase coding sequence) restriction sites. We subsequently reconstructed the expression cassette using four PCR products:

1) The Gal1 promoter delta Gal4 with 8 operator LexA and the Kozack sequence with a new downstream MCS (*Asc*I, *Nhe*I) (primSB_0076 and 0077).
2) The Gal1 Nterm sequence (I10-C20), originally expressed by the pSH18-34, is used as spacer (primSB-0078 and 0079) between the two copies of the Tag BFP.
3) The coding sequence of the Tag-BFP (from pTag_BFP-Actin, Evrogen) borded with 2 *Xho*I sites, (primSB_0084 and 0085).
4) The terminator sequence (primSB_0080 and 0081).

These 4 amplicons were then used to perform directly a gap repair in yeasts. Thus, we obtained the pSB_3RO plasmid. A second copy of the Tag-BFP (primSB_0120 and 121) was inserted in our new *Nhe*I site (Thermo Scientific), by homologous recombination to generate the pBFP2 plasmid. This final vector allows the expression of a dimer of Tag-BFP as reporter of the yeast two hybrid reaction.

### Western blot

Total protein extracts were obtained from 6 OD_590nm_ exponentially growing diploids yeasts as previously described (32) into 60µl of sample buffer. Ten microliters were used for SDS-Page analysis on Bolt™ 4-12% Bis-Tris Plus Gels (Thermo Scientific). Electrophoresis separation was performed in NuPAGE™ MOPS SDS Running Buffer (Thermo Scientific). Proteins were then transferred on a Nitrocellulose Membrane 0.45µm (Biorad), using a Trans-Blot® Turbo™ Transfer System (Biorad) for 14 minutes, at 1A and 25V. The membrane was subsequently blocked 1 hour at room temperature in PBS + tween 0.2% (v/v) supplemented with 5% (w/v) low-fat milk powder. HA tagged proteins were labeled overnight at 4°C with the mouse HA.11 Clone 16B12 Monoclonal Antibody (Eurogentec) 1/2000 in PBS + tween 0.2% (v/v) + 10 mg/ml BSA (Albumin bovine fraction V, Euromedex). The membrane was then washed four times seven minutes in blocking buffer at room temperature. The membrane was then incubated for one hour at room temperature in presence of a sheep anti-mouse whole IgG HRP (GE Healthcare) secondary antibody diluted 1/5000 in blocking buffer. The excess of antibody was removed with two washing steps of five minutes in PBSt at room temperature. Labelled proteins were then revealed with Super Signal West Pico chemiluminescent substrate (Thermo Scientific) using a Biorad Chemidoc apparatus, following instructions provided by the suppliers.

### qY2H in liquid phase

Chemo-competent EGY42 (MATa; trp1, his3, ura3, leu2) and TB50 (MATα; trp1, his3, ura3, leu2, rme1) yeasts were generated as previously described (33).

Competent EGY42a yeasts were transformed with 1µg of pBFP2 and grown on selective SD-U medium. Chemo-competent EGY42a pBFP2 yeasts were then generated and transformed with 1µg of Bait vectors. Haploid Bait yeast strains were then selected on SD-UH medium. Competent TB50α yeasts were transformed with 1µg of Prey vector. Haploid Prey yeast strains were selected on SD-W medium. Matrix mating assay were performed for one night with 50µl of Bait and Prey strains (each) resuspended in YPAD medium at 0.1 OD at 30°C. The next morning YPAD medium was removed and the yeast diploids were harvested and amplified in 1ml of SD- UHW for 3 days at 30°C.

The qY2H assay was performed in pre-heated (30°C) and oxygenated SGR-UHW supplemented with Galactose 0.25% (Euromedex) and Raffinose 1% (Sigma-Aldrich) to induce the expression of the Prey proteins. To ensure we obtained an excessive number of cells (about 10^7^) for the analysis, a culture of 100 ml was inseminated with 600 µl of saturated diploids per couple of interest. It turned out that for a typical analysis a number of 10^6^ cells is adequate, so that actually 10 times smaller cultures and insemination volumes can be used. The yeasts were incubated for 2h at 30°C without shaking, and then harvested after a centrifugation step of 10min at 1000g. The yeast were resuspended in 1ml PBS (Dominique Dutscher), centrifuged 1min at 13000 rpm, and washed again with 1ml of PBS. The yeasts were resuspended in 500µl of PBS 4% PFA (Sigma-Aldrich, Catalog n°P6148) and incubated for 10min at 4°C. The fixation reaction was blocked by 2 washing steps with 1ml PBS, and one incubation of 15min at 4°C in 500µl of PBS 0.1M Glycine (Euromedex). Finally, the yeasts were washed twice in PBS, and stored in 1ml of PBS at 4°C for not longer than 24 hours.

### Flow cytometry and data analyses

The expression levels of BD-Bait, AD-Prey and reporter were acquired in linear scale using a MacsQuantVYB flow cytometer (the settings are presented in Suppl. Table S2). To ensure homogeneous sampling of the yeasts cells in suspension, we used the strong mixing mode. With the apparatus at our disposal, this mode generates at very early acquisition times a small population of particles with abnormal characteristics for yeast cells (a high red fluorescence intensity, even for non-fluorescent samples). We suspect these are micro-bubbles. To rigorously eliminate this population, we skipped the first 20 000 events of all samples files in the subsequent analysis. The flow-cytometry files were analyzed using the FlowCytometryTools package for python (http://eyurtsev.github.io/FlowCytometryTools). When a hlog-transformation (34) was applied, the following settings were used: b = 2000, r = 10000, and d = 5.42.

Our python scripts (with a graphical user interface and user guide for the automated generation of the qY2H affinity ladder) can be downloaded here: http://github.com/LBMC/qY2H-Affinity-Ladder. The flow cytometry files of the experiment shown in Fig. 4 can be downloaded from http://flowrepository.org under accession number FR-FCM-ZYUL.

## Results

### qY2H enables the monitoring of the expression level of the reaction partners after 2h

Two hours after induction of the Y2H reaction, the expression of all BD-Bait and AD-Prey fusions can be easily detected. Fig. 2A displays their fluorescence intensities for a subset of couples. The fluorescence intensity typically spans several orders of magnitude higher than the negative controls. However, expression problems can be seen for BD-HRas (Fig. 2C). Independent quantification by Western Blotting (Fig. 3) shows that the expression level of BD-HRas is indeed impaired. In fact, BD-HRas cannot be detected in the Western blot. Flow cytometry, on the other hand, indicates a slight shift in the probability distribution for BD-HRas expressing cells (with respect to the negative control, see also Fig. 4A).

**Fig. 2.**
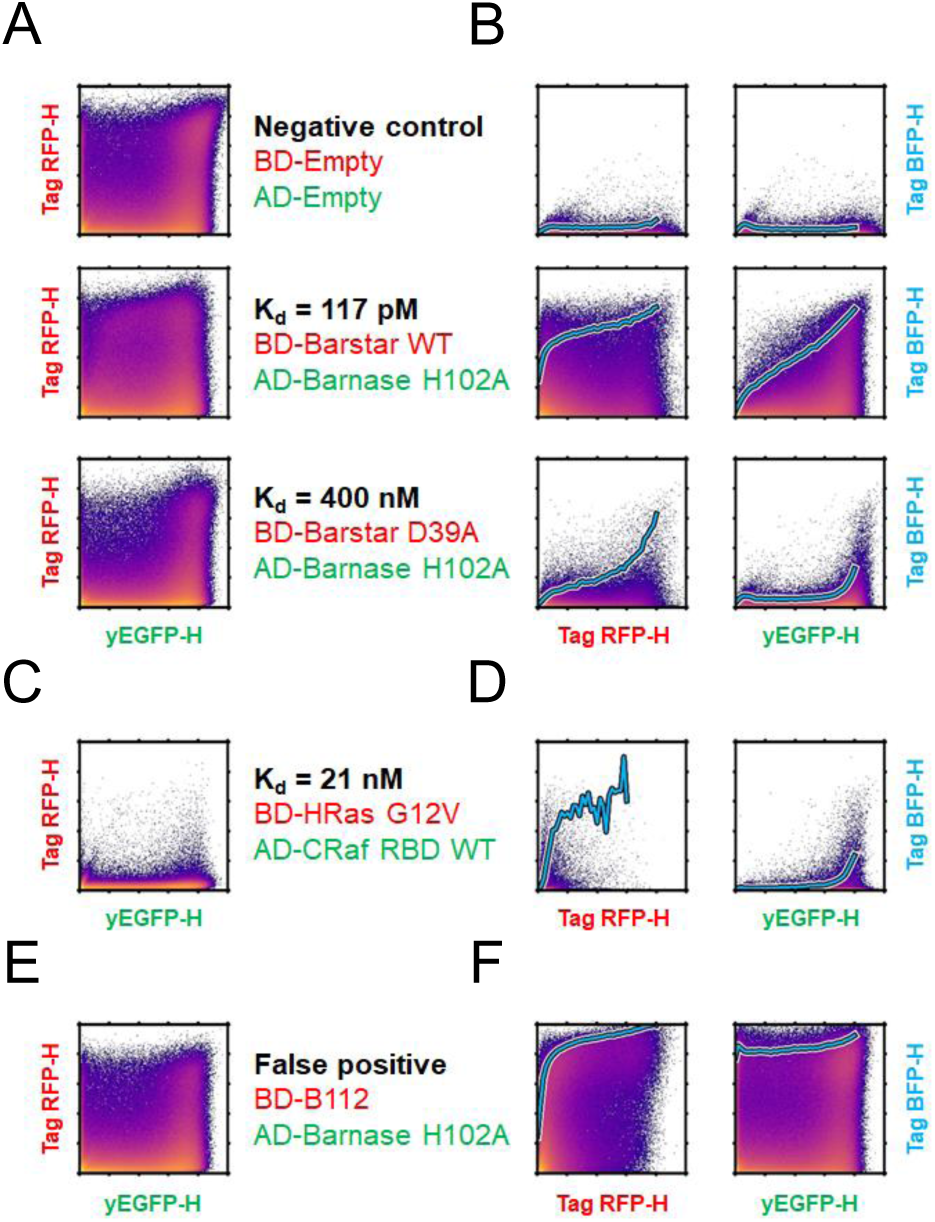
Single-cell raw data from the flow-cytometry acquisition. *A-F*. Fluorescence intensities in the channels corresponding to the BD-Bait (Tag-RFP-H), AD-Prey (yEGFP-H) and reporter (Tag-BFB-H) are represented as density-colored scatter plots for a subset of studied couples. Intensities were pre-processed by a hlog-transformation. The expression of the BD-Bait as a function of AD- Prey is presented in *A*, *C* and *E*. The expression level of the reporter as function of the BD-Bait or AD-Prey is displayed in *B*, *D* and *F*. The cyan bold line represents the evolution of the Tag-BFP- H intensity when averaged over the 5% top-ranked cells in slices of the yEGFP-H or Tag-RFP-H channel. Our system was validated using a negative control, a strong (picomolar) and a medium (nanomolar) couple, see *A* and *B*. As an example for low expression, the construct BD-Bait HRas is shown in *C* and *D*. We also constructed a genetic fusion of the LexA BD with the B112 AD as an example of a false positive interaction, see *E* and *F*.

**Fig. 3.**
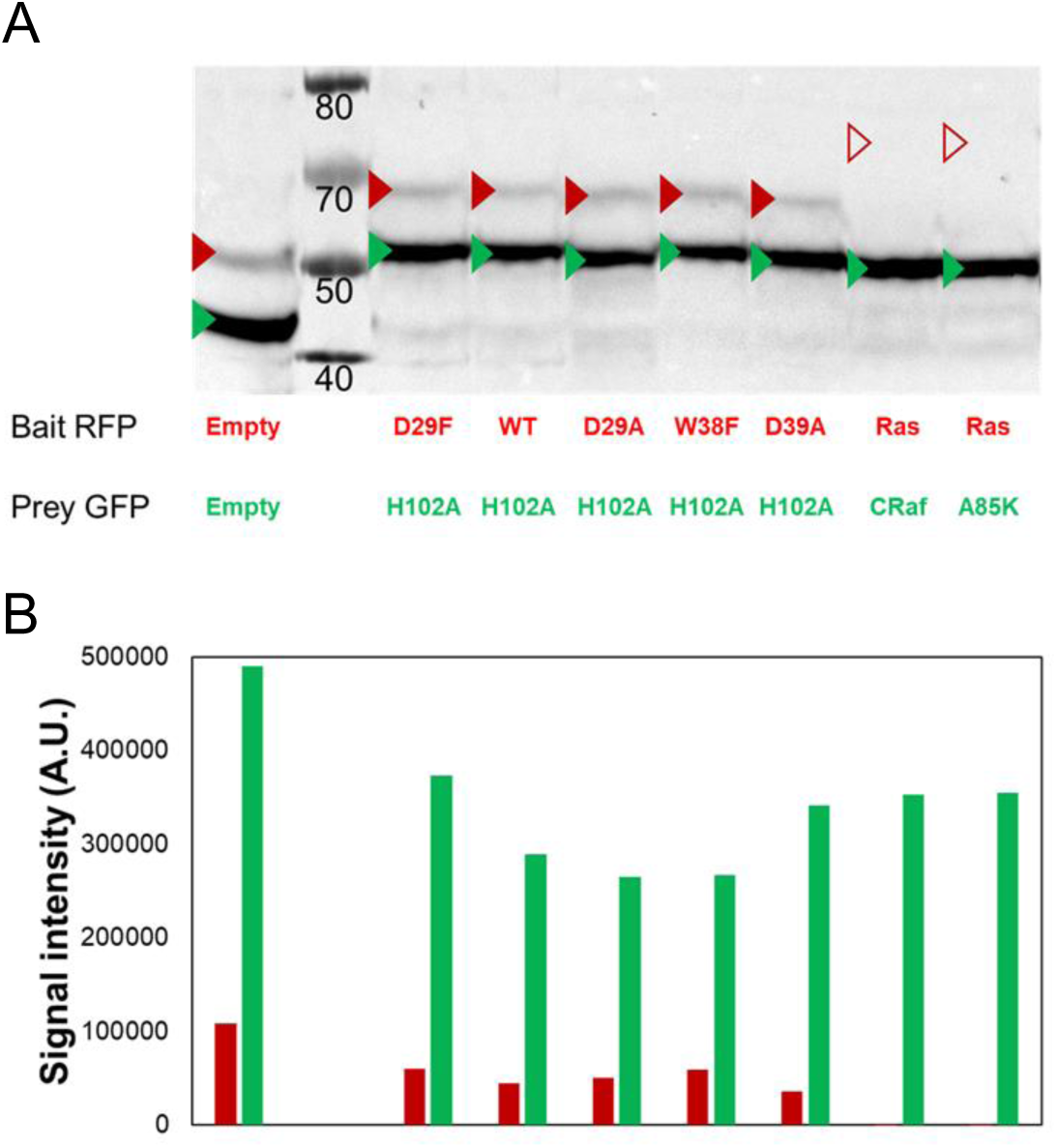
Simultaneous detection of BD-Baits and AD-Preys by western blotting using a single primary antibody. *A*. The various studied couples were submitted to western blot analysis. Total proteins were extracted from 6 OD of diploid yeasts grown for 2h at 30°C in SGR –UHW (0.25% galactose). The BD-Bait and AD-Prey fusion proteins were simultaneously detected by Western blotting using HA tag (originally present only in the AD-Prey proteins, and newly added to the BD-Bait proteins). The expected molecular weights of the fusion proteins are indicated by red (BD-Bait) or green (BD-Prey) triangles. Except for Ras G12V C186A (empty red triangles), all the proteins are detectable at their correct molecular weight. *B*. The differential expression level was then quantified using ImageJ program. A four to nine fold overexpression of the AD-Prey compared to the corresponding BD-Bait (when detectable) can be observed.

**Fig. 4.**
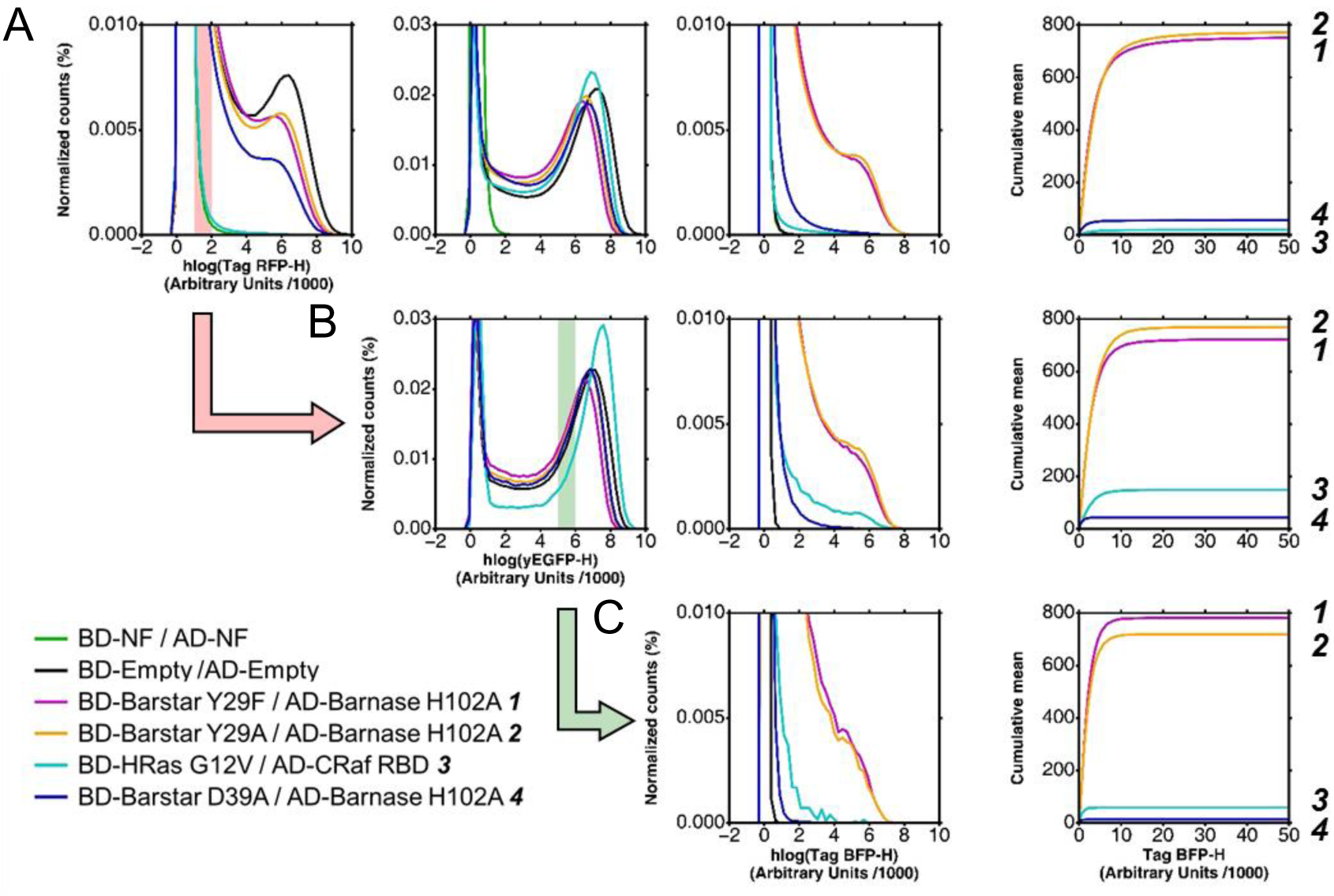
Influence of the expression level of BD-Bait and AD-Prey on the read-out. *A-C*. The expression levels of BD-Bait (1^st^ column), AD-Prey (2^nd^) and reporter (3^rd^) are visualized as distribution of the hlog-transformed fluorescence intensities. In order to better visualize differences in the reporter expression we use an additional representation of the cumulative mean in linear scale (4^th^ column). The distributions and cumulative mean are shown for the global population in *A* and gated subpopulations in *B* and *C*. For reasons of clarity only an illustrative subset of studied couples is shown. In the legend these couples are ordered and numbered (***1***-***4***) according to their *in-vitro* affinity. The order of the couples according to the reporter expression (*i.e.*, mean value of the Tag-BFP-H channel) is given on the very right side of the three subfigures. The global population of cells reveals significant differences in the expression levels of BD-Bait and to a smaller extent of AD-Prey *A*. Due to these expression level differences, the mean values of Tag-BFP-H are ordered differently than the *in-vitro* affinities: ***2*** displays a stronger reporter expression than ***1***, and ***4*** stronger than ***3***. Only the discrimination between strong (***1***, ***2***) and medium (***3***, ***4***) interactors is possible. The successive gating of cells, first based on Tag-RFP-H values (*B*), and then based on yEGFP-H values (*C*), allows to equalize the expression levels. Such standardized subpopulations improve the extraction of quantitative information on the strength of the Bait-Prey interactions: the Tag-BFP-H mean value is ordered according to the *in vitro* affinities. The transparent red bar in the panel of the 1^st^ column shows the Tag-RFP-H gating interval applied to obtain the subpopulation in *B* (symbolized by the red action arrow). Similarly, the green bar and arrow illustrate the yEGFP-H gating interval applied to the subpopulation in *B* to obtain the final subpopulation in *C*.

Thus, flow cytometry gives an immediate indication on eventual expression problems of the Bait and Prey fusions during acquisition. This contrasts with standard Y2H experiments where the expression level of the BD-Bait and AD-Prey fusions is usually unknown at the time of reporter detection. More laborious Western Blots are usually required to gain this information.

### Reporter level is correlated with the expression level of the reaction partners

Even for the weakest Bait-Prey interaction the reporter can already be detected two hours after induction (Fig. 2); it is clearly above the level of the negative control (panel B). Moreover, we observe that the reporter level (*i.e.*, blue fluorescence intensity) is correlated with the green and red fluorescence intensity: more reaction partners (*i.e.*, higher amount of interacting BD-Bait and AD-Prey fusions) yielded more product (reporter). This obvious correlation has consequences for the extraction of quantitative information on the strength of PPIs as we demonstrate later on. In the case of the Bait/Prey-couple BD-B112/AD-Barnase H102A (Fig. 2F) this correlation is basically only observed for the red fluorescence (BD-Bait). The B112 acid blob acts as an activation domain (35, 36) so that this specific BD-Bait fusion is a functional transcription factor by itself that does not depend on the AD-Prey fusion.

### Standardization of BD-Bait and AD-Prey levels is required to gain information on binding strength

In all repetitions of the experiment, the reporter level of the global cell population roughly reflects the magnitude of the (*in-vitro*) affinity. As shown in Fig. 4A for a single experiment and a small subset of PPIs, the Bait-Prey couples with high affinity (K_d_ ∼ pM) could be easily distinguished from medium-affinity couples (K_d_ ∼ nM) based on their mean reporter level. It confirms results of previous studies that the global Y2H read-out correlates with *in-vitro* affinity (13–17). In addition, our approach discloses the influence of the expression levels of the reaction partners, *i.e.*, the BD-Bait and AD-Prey fusions. Their levels may vary significantly between the studied couples (Fig. 2 & 3) and to a smaller extent between different experiments. These variations complicate the discrimination of Bait-Prey couples based on their affinities. Fig. 4A presents a particularly illustrative experiment: The couple BD-HRas / AD-CRaf displays a 20 times lower *in-vitro K_d_*-value than the couple BD-Barstar D39A / AD-Barnase H102A. Yet the mean reporter level (Fig. 4A, right column) is lower for the former couple than for the latter couple; the opposite would have been expected according to the *in-vitro* affinities. We note, however, that the former couple exhibits a significant lower expression level of the BD-Bait fusion (Fig. 4A, left column) than the latter. Thus, can quantitative information on the strength of the interaction (*i.e.*, relative affinities) be reliably extracted from such an experiment?

To address this question, we tried to correct for differences in the expression level by sub-selecting (gating) only cells that display a red and green fluorescence intensity within a certain narrow interval (see Fig. 4B & C). And indeed, when standardizing the red fluorescence intensity (*i.e.*, gating for cells with similar BD-Bait expression level), the mean reporter level reflects the *in-vitro* affinities for the two couples BD-HRas / AD-CRaf and BD-Barstar D39A / AD-Barnase H102A (Fig. 4B). Similarly, the couple BD-Barstar Y29F / AD-Barnase H102A has the lowest AD-Prey expression level and displays a weaker reporter level than expected. Standardization of the AD-Prey level corrects the reporter levels of the studied couples according to their reported *in-vitro* affinities (Fig. 4C). Changing the location of the gating intervals leads to the same conclusions (see Suppl. Fig. S2). Nevertheless, suggestions can be made to choose the optimal the gating intervals (see “Recommendations”).

### Affinity ladder permits rapid classification of PPIs based on their strength

Often the goal is to rank PPIs based on their affinity or to obtain an upper and lower bound for the dissociation constant. With the above gating approach, an affinity ladder can easily be generated with a set of PPIs with known dissociation constants (Fig. 5). Standard software of flow cytometers can be used to perform the required gatings and calculate mean fluorescence intensities. We provide a graphical user interface to automate the generation of the affinity ladder (see Experimental Procedures).

**Fig. 5.**
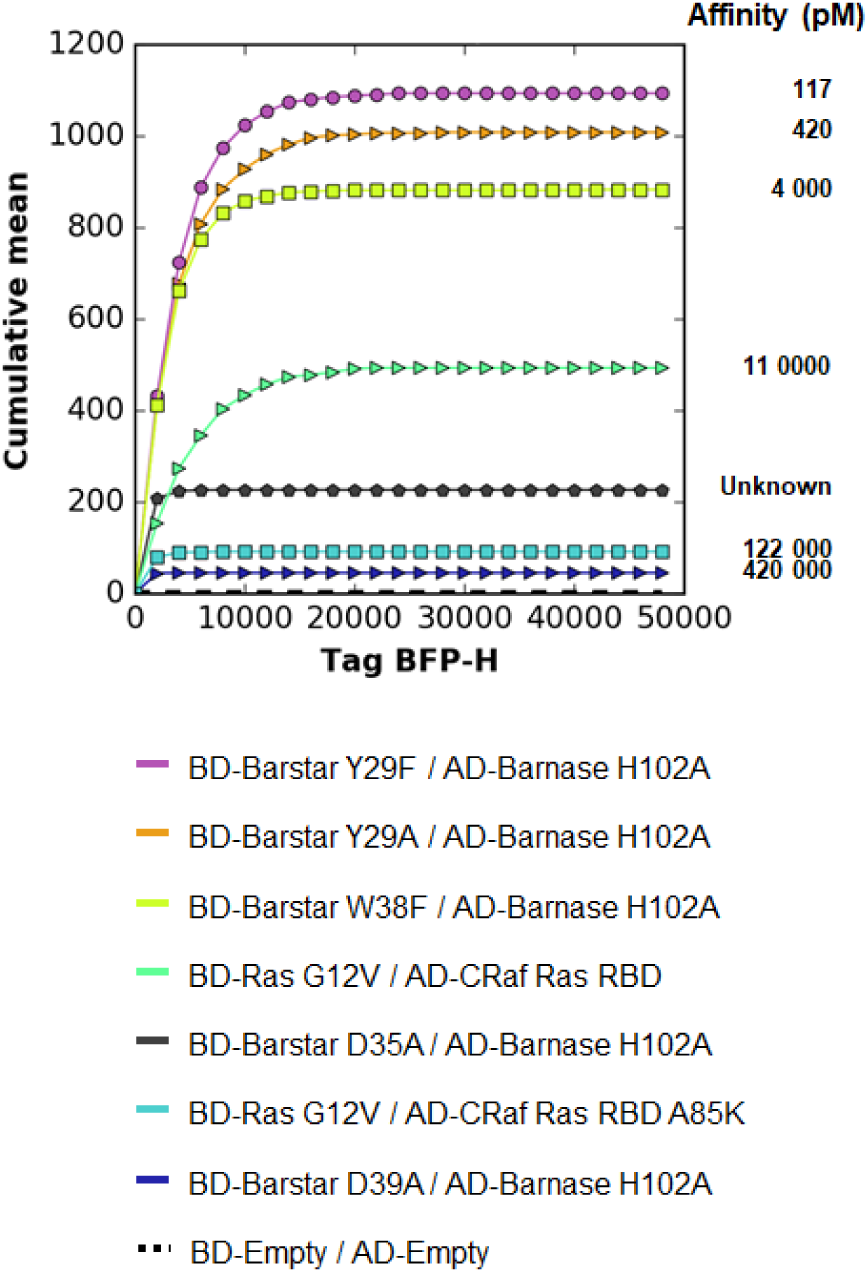
qY2H affinity ladder. The same dual gating approach as in Fig. 4 was applied to all studied couples of Table 1. Here one representative experiment is presented. The cumulative mean of the negative control (BD-Empty / AD-Empty) was removed in all cases. The resulting cumulative mean curves are ordered according to their reporter dissociation constants (Table 1). For the couple BD-Barstar D35A / AD-Barnase H102A no dissociation constant is reported in the literature. Our affinity ladder allows to rank the constant between 11 and 122 nM.

The generated affinity ladder can then be used for a rapid visual classification of PPIs with thus far unknown affinities within a given range (here from nano- to picomolar). This is demonstrated at the example of the mutation D35A of Barstar. Thus far, no *in-vitro* affinity data is available for the interaction of this mutant with Barnase H102A. With the affinity ladder of Fig. 5 we can rank the affinity between those of HRas/CRaf 122 nM) and HRas/CRafA85K (11 nM).

Our experiment indicates that the Barstar mutant D35A exhibits a significantly higher affinity for Barnase H102A than the Barstar mutant D39A (420 nM). To validate this observation, we performed independent alchemical free-energy calculations (see Supp. Mat.). Through the use of a thermodynamic cycle (Suppl. Fig. S3A) we calculated the difference in binding free energy between the mutants Barstar D35A and Barstar D39A. We obtained a value of −1.9 ± 0.3 kcal/mol which indicates that the dissociation constant of the mutant D35A is about 20 times lower than that of the mutant D39A. Thus, we can estimate a dissociation constant of about 20 nM for the mutant D35A in agreement with the qY2H experiment.

## Discussion

### Quantitative features of the tri-fluorescent yeast two-hybrid system

The tri-fluorescent qY2H system offers the novelty of identifying expression correlations for the genes involved in the actual Y2H reaction. For true positive interactions the read-out is correlated with both reaction partners. For false positive interactions where the Bait acts as an AD (*e.g.*, B112), the read-out is basically only correlated with one reaction partner. This characteristic correlation patterns can serve as additional criteria to discriminate such false positive interactions from true positives. It complements the use of proper controls (*i.e.*, empty Bait and Prey plasmids) routinely applied in Y2H assays.

In the past, significant effort has been spent to render the Y2H read-out quantitative and thereby gain quantitative information on the strength of interactions (see cited literature in the Introduction). Our study clearly demonstrates that the quantification of the reaction partners is important, too. We have shown that variations in the expression levels of BD-Bait and AD-Prey can lead to reporter levels that are not ordered according to the underlying PPI affinities. Through a simple gating process, it is, however, possible to standardize the expression levels of BD-Bait and AD-Prey and thereby overcome this difficulty.

### In-cellula is not in-vitro

The observed agreement between *in-cellula* reporter levels and *in-vitro* affinities (Fig. 4B) cannot be presumed *a priori*. The *in-vitro* experiment measures the affinity between the interactors alone (or with tags) whereas the qY2H system relies on fusions proteins (Fig. 1). If the fused domains influence the interaction between the Bait and Prey, *e.g.*, by blocking the binding interface, the resulting *in-cellula* reporter level would be impaired and most likely not correlate with the *in-vitro* affinity.

*In-vitro* experiments measure the affinity under well-defined buffer-controlled equilibrium conditions. In contrast, our *in-cellula* experiments take place in non-equilibrium microvessels (37) where the interaction partners can interact with the endogenous complex solution of biomolecules. This may lead to effectively smaller concentrations of the reaction partners. Also, post-translational modification(s) could impact the interactions.

With prior knowledge about the Bait or Prey the amino-acid sequence can be optimized to take into account some effects, like specific sub-cellular localization. An illustrative example is the protein HRas, which is usually found to be associated to the cytoplasmic membrane through its C-terminal anchor. Mutation of C186A abolishes the anchor function (38) and the protein can be used for the *in-cellula* titrations.

### Recommendations

Beside potential sequence optimizations (as proposed above), we recommend the following precautions to be taken for the measurement with the qY2H system:

1) As in any Y2H screen, BD-Bait and AD-Prey constructs should be tested against negative controls, *i.e.*, an empty AD and BD construct, respectively, to identify potential false positive interactions. Also, a mating of yeasts expressing only empty (but fluorescent) AD and BD constructs is recommended; it serves to remove the background of the system.
2) The PPIs should be tested in both orientations, *i.e.*, with the proteins switched between the Bait and Prey vectors, to identify the orientation with the lower background and with the higher reporter level (for standardized levels of reaction partners).
3) We recommend to pre-transform BD-Bait-expressing haploids with the read-out plasmid; it increases the read-out; two subsequent transfections are more efficient than a single double transfection. Use only freshly transformed yeast cells for the qY2H experiment. Storing diploids yeast cells for a week in the refrigerator decreases the level of AD-Prey and read-out by a factor two to five.
4) For the construction of the affinity ladder, the gating interval for the red fluorescence intensity (BD-Bait) was positioned at the lowest possible location to avoid saturation effects, *i.e.*, it was set just above the 95-% threshold of the negative control. The gating intervals of the green fluorescence intensity was set to a medium range value to reach the desired sensitivity but to avoid saturation and protein burden effects (39). The width of each interval gate should be about 20-30% of the value of its lower border.
5) If the gating intervals are not directly applied at acquisition time on the flow cytometer, at least 10^6^ cells should be acquired for analysis. This number is sufficient to reach a converged ladder after gating (see Suppl. Fig. S4). For this number of cells cultures of 10ml are sufficient.

## Conclusion

The newly constructed vectors provide access to a novel quantitative Y2H system with fluorescent tags for the reaction partners (BD-Bait, AD-Prey) and the reporter. The established protocol is rapid, sensitive and highly reproducible. It permits easy detection of expression problems of the reaction partners. Using flow cytometry, the expression levels of the reaction partners can be monitored cell by cell simultaneously with the level of the reporter. The single-cell data can be exploited to identify correlation patterns as indicators of physical interactions.

The qY2H method presented in this work offers also an approach to quantitative data on the strength of protein-protein interactions in living cells. In this context, we have demonstrated the importance of quantifying the product <u>and</u> the reaction partners of the Y2H reaction: standardization is critical to correct for differences in expression levels between couples. Using a straightforward gating analysis, an affinity ladder can be easily generated that permits rapid classification of PPIs according to their affinity. We would like to emphasize, however, that these *in-cellula* affinities are effective quantities that depend on the cell’s complex microenvironment; and this environment may change as a function of the yeast strain and the experimental conditions (temperature, medium, *etc*).

All steps of the protocol have been optimized in liquid phase that can be easily automated for the use of microplates and integrated within robotic pipelines. It sets the stage for high-throughput quantitative Y2H screenings of PPIs using cross-mating approaches (40, 41) with libraries of yeast clones. As an outlook, quantitative PPI networks can be created by attributing weights to the PPI edges according to their *in-cellula* reporter levels. It contrasts standard Y2H screens that yield networks with only binary information (YES or NO). The topology of force-directed networks may help identifying key pathways within the network, and how these paths change as a function of environmental conditions (stress, metabolism, *etc*). Thus, we anticipate that high-throughput qY2H data would boost the modelling of interactomes and thereby advance significantly systems biology.

## Acknowledgements

This project was supported by a grant from the Fond Recherche of the ENS de Lyon. We are grateful to the Pôle Scientifique de Modélisation Numérique (Lyon, France) for computer time, and the SFR Biosciences Gerland-Lyon Sud (UMS344/US8) for the access to the MacsQuantVYB flow cytometer. We thank Dr Francesca Palladino and Matthieu Caron for technical support on the Western blots and Dr Gaël Yvert for helpful comment on the manuscript.

## Data Availability

Our python scripts (with a graphical user interface and user guide for the automated generation of the qY2H affinity ladder) can be downloaded here: http://github.com/LBMC/qY2H-Affinity-Ladder.

The flow cytometry files of the experiment shown in Fig. 4 can be downloaded from http://flowrepository.org under accession number FR-FCM-ZYUL. Other experimental data can be provided upon request.

## Abbreviations

PPIs: Protein-Protein interactions
qY2H: quantitative yeast two-hybrid
BD-Bait: DNA Binding Domain fused to the Bait
AD-Prey: Activation Domain fused to the Prey

