## Supplementary Material for "A quantitative tri-fluorescent yeast two-hybrid system: from flow cytometry to *in-cellula* affinities"

David Cluet, Ikram Amri, Blandine Vergier, Jérémie Léault, Clémence Grosjean, Dylan Calabresi, and Martin Spichty[#]

Laboratoire de Biologie et de Modélisation de la Cellule, Ecole Normale Supérieure de Lyon, CNRS, Université Lyon 1, Université de Lyon; 46 allée d'Italie; 69364 Lyon cedex 07; France.

Running title: **A quantitative tri-fluorescent yeast two-hybrid system**

### Computational methods for the alchemical free energy calculations

We used a standard dual topology approach where the two residues 35 and 39 of Barstar were simultaneously transformed from A to D and from D to A, respectively, as a function of the usual coupling parameter λ. The reader is referred to the literature for more details (1).

*Software*

Model systems were set up with the program CHARMM (2), version c39b1. Initial input files for CHARMM were generated with the CHARMM-GUI server (3) and then modified to implement the actual structural model with dual topology (see below). Molecular dynamics simulations for free energy calculations were carried out with the program NAMD, version 2.11 (4).

*Model system*

The crystal structure of the complex Barnase:Barstar (5) served as starting point of the structural model. For the alchemical transformation in Barstar alone (see left, vertical leg in Suppl. Fig. S3A) only the coordinates of chain D, residues 1-89 (PDB entry 1BRS), were selected. For the transformation of the complex (right vertical leg) the coordinates of Barnase (chain C, residues 1-110) were specified in addition to those of Barstar. Missing residue coordinates were added with the help of the CHARMM-GUI server. Both systems were solvated in a cubic box (side length of 80 Å) of TIP3 water molecules. With the program CHARMM, residues 35 and 39 were then replaced by residues with dual topology for aspartic acid (D) and alanine (A).

*Settings for molecular dynamics*

The systems were simulated with the CHARMM36 force field and periodic boundary conditions. A cutoff of 12 Å was used for short range non-bonded interactions whereas long-range electrostatic interactions were treated by Particle-Mesh Ewald (PME) with a grid spacing of 1 Å. The equation of motion was integrated with a time step of 1 fs. Short range non-bonded forces were updated every 2 fs, PME forces every 4 fs. The system was kept at constant pressure (1 atm) and temperature (298.15 K) with a piston oscillation period of 100 fs and a damping time scale of 100 fs. With these MD settings the average box size of both systems (Barstar alone and complexed with Barnase H102A) was almost identical (77.77 ± 0.05 vs. 77.64 ± 0.05 Å) in the following free energy calculations. For systems with identical (cubic) box sizes and no change in net charge (during the alchemical transformation), finite-size effects should largely cancel when comparing the two legs of the alchemical transformations (6–8).

*Alchemical transformations*

The electrostatic interactions of the outgoing residues (here A35, D39) are linearly removed from $\lambda$=0 to $\lambda$=0.5 and those of the incoming residues (D35, A39) linearly added from $\lambda$=0.5 to $\lambda$=1 (NAMD parameter alchElecLambdaStart=0.5). The van-der-Waals interactions are scaled down from $\lambda$=0 to $\lambda$=1 for the outgoing residues, and scaled up from $\lambda$=0 to $\lambda$=1 for the incoming residues (alchVdWLambdaEnd=1). To avoid endpoint problems, we used NAMD's soft-core potentials with a shifting coefficient of 4 Å. Non-bonded interactions within outgoing and incoming atoms were also scaled with $\lambda$ to account for interactions between residues 35 and 39 (alchDecouple=off). The actual transformation was done with a windowing method where the dimer and tetramer systems were sampled by molecular-dynamics at 21 equally-separated $\lambda$-values (from 0 to 1 with an increment parameter of 0.05). For each window we performed two separate MD simulations. Using NAMD's FEP option, we recorded on-the-fly every 1000 fs the work required to switch instantaneously the Hamiltonian from the actual window to one of the neighboring windows. For the endpoints at $\lambda$=0 and 1 we performed only one MD simulation (because there is only a single neighbor).

The free energy difference between two neighboring windows was calculated from the collected work data for forward and reverse switches using Bennett's acceptance ratio (BAR) method (9). The total free energy change for the transformation of $\lambda$ from 0 to 1, (= $\Delta G_{\mathrm{alchemical}}$, see also Suppl. Fig. 3A) was obtained by summing up all differences between neighboring windows. After an equilibration phase of 1 ns (Barstar alone) and 5 ns (complex), we monitored for blocks of 250 ps (Barstar) and 1 ns (complex) the value of $\Delta G_{\mathrm{alchemical}}$ (Suppl. Fig. S3B). The value of $\Delta G_{\mathrm{alchemical}}$ fluctuates around a mean value of -0.17 ± 0.08 kcal/mol in the case of the Barstar alone and 1.71 ± 0.31 kcal/mol in the case of the Barstar-Barnase complex. The errors correspond to twice the standard error of the mean. Finally, the difference between the two vertical legs of the thermodynamic cycle of Suppl. Fig. S3A can be calculated:

$$\Delta G^{\mathrm{Barstar}}_{\mathrm{alchemical}} - \Delta G^{\mathrm{complex}}_{\mathrm{alchemical}} = -1.88 \pm 0.32\ \mathrm{kcal/mol}$$

The usual error propagation rule was used for this subtraction.

**Suppl. Table S1: Primers list.**

| Primer name | Sequence |
|---|---|
| primSB_0001 | GTTGGGGTTATTCGCAACGGCGACT |
| primSB_0002 | GAAATTCGCCCGGAATTAGCTTGGCT |
| primSB_0003 | CCTTATGATGTGCCAGATTATGCCTCTCCCGAATTCATGGTGTCTAAGGGCGAAGAGCTGATT |
| primSB_0004 | GACTGCTTTTTTCATCTCGAGGGCGCGCCCGAATTCATTAAGTTTGTGCCCCAGTTTGCTAGGG |
| primSB_0018 | GGGCACAAACTTAATGAATTCGGGCG |
| primSB_0019 | AGCTTGGCTGCAGGTCGACTCACT |
| primSB_0010 | CCTTATGATGTGCCAGATTATGCCTCTC |
| primSB_0011 | CCAAACCTCTGGCGAAGAAGTCCAAA |
| primSB_0012 | CCTTATGATGTGCCAGATTATGCCTCTCCCGAATTCATGAGCAAGGGCGAGGAGCTGTTC |
| primSB_0013 | GTTGATAACCTGTGCCTCGAGGGCGCGCCCGAATTCCTTGTACAGCTCGTCCATGCCGAG |
| primSB_0020 | GACGAGCTGTACAAGGAATTCGGG |
| primSB_0021 | AAGTCCAAAGCTTCCATGGTCACTCACTTA |
| primSB_0076 | GGACGCAAAGAAGTTTAATAATCATATTACATGGC |
| primSB_0077 | GAAAAAACTATAATGACTAAATCTCATTCAGAAGAAGTGGGGCGCGCCGCTAGC |
| primSB_0078 | CATTCAGAAGAAGTGGGGCGCGCCGCTAGCATTGTACCTGAGTTCAATTCTAGCGCAAAGG |
| primSB_0079 | CCAAGCTTGGCCAAGCCCGGACTCGAG |
| primSB_0080 | TAACTCGAGTAATAACCGGGCAGGCCATGTCTG |
| primSB_0081 | TAATAAAACGCCCGTTCCCGGACG |
| primSB_0084 | ATTCCAAGCTTGGCCAAGCCCGGACTCGAGATGAGCGAGCTGATTAAGGAGAACATGC |
| primSB_0085 | CCTAGCAAACTGGGGCACAAGCTTAATTAACTCGAGTAATAACCGGGCAGGCCATGTCTG |
| primSB_0120 | AAATCTCATTCAGAAGAAGTGGGGCGCGCCGCTAGCATGAGCGAGCTGATTAAGGAGAACATGC |
| primSB_0121 | CTTTGCGCTAGAATTGAACTCAGGTACAATGCTAGCATTAAGCTTGTGCCCCAGTTTGCTAGG |

**Suppl. Table S2: Macsquant VYB settings for qY2H fluorescence acquisition.**

| Channel | Setting |
|---|---|
| FSC | 229V |
| SSC | 265V |
| V1 | 264V |
| B1 | 327V |
| Y1 | 506V |

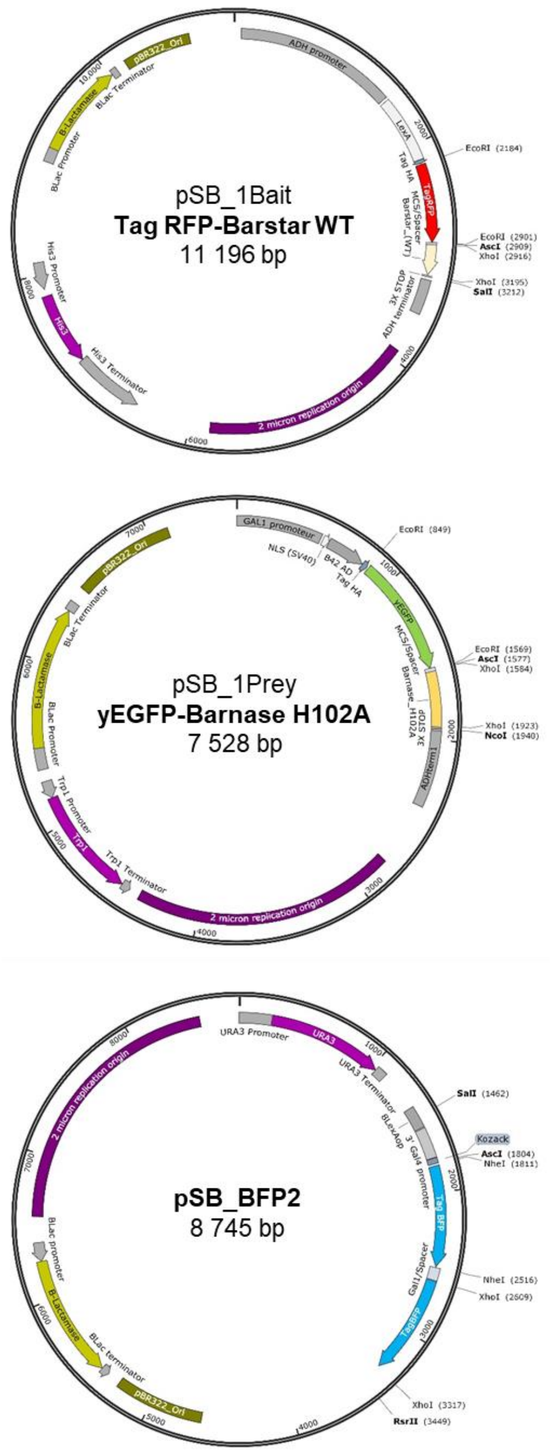


**Suppl. Fig. S1.** qY2H plasmid maps.

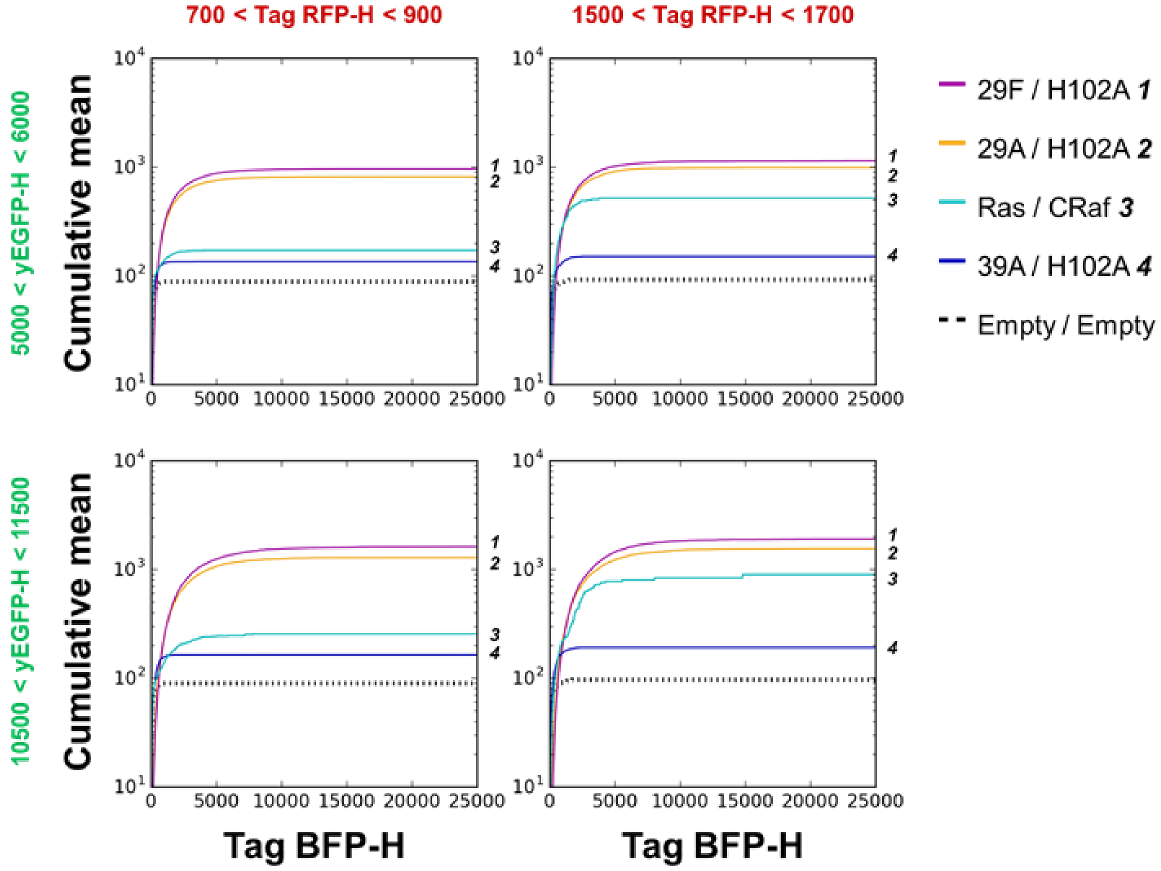


**Suppl. Fig. S2.**

**Impact of the gating region on the qY2H affinity ladder.** The same dual gating approach as in Fig. 4 was used with two different intervals for each channels: 700-900 and 1500-1700 for Tag RFP-H, and 5000-6000 and 10500-11500 for yEGFP-H. The four different combinations do not affect the ordering of the couples. Only their relative positions vary from one gates combination to another.

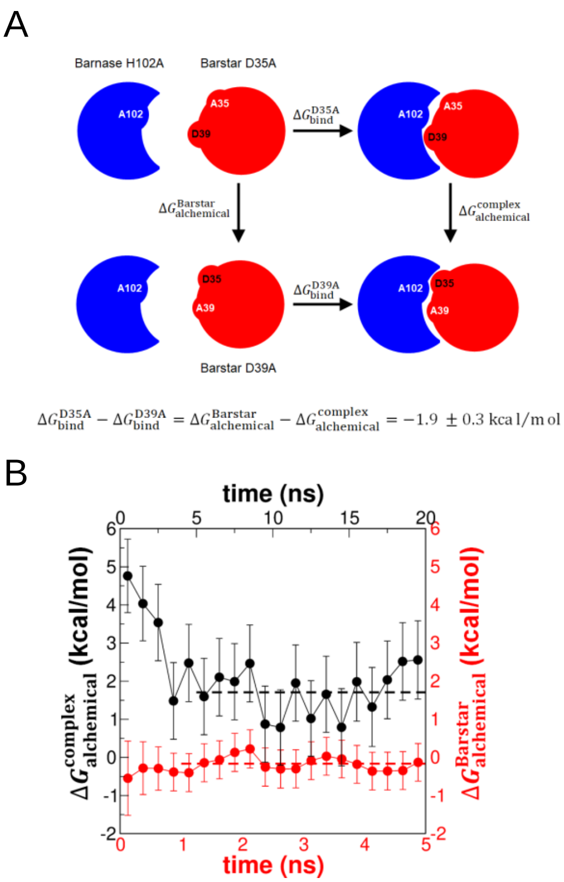


**Suppl. Fig. S3.**

**Alchemical free energy calculations**. *A.* A thermodynamic cycle is applied to calculate the difference in binding free energy for the interaction between Barnase H102A and the mutants Barstar D35A (horizontal leg, top) and D39A (horizontal leg, bottom). Because the free energy is a state function, this difference can also be obtained from the free energy difference of the corresponding alchemical transformation in Barstar alone (vertical leg, left) and the complex (vertical leg, right leg). *B.* In the block analysis the change in free energy for the alchemical transformations is plotted for consecutive blocks of 250 ps (Bartsar alone, red dots) and 1 ns (complex, black dots) of sampling. The error bars correspond to the analytical error of the maximum likelihood estimate (10). The mean value of $\Delta G_{\mathrm{alchemical}}$ for each alchemical transformation is indicated as horizontal dashed line.

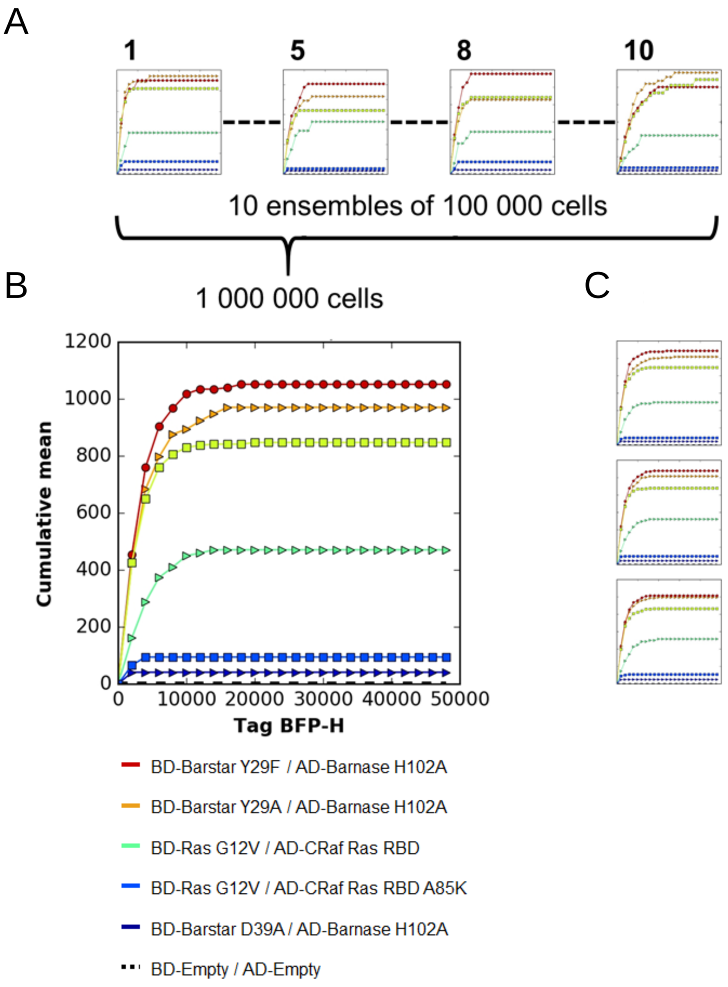


**Suppl. Fig. S4.**

**Impact of the number of cells on the qY2H affinity ladder.** *A*. Ten successive sub-ensembles of 100 000 cells from one single experiment were used to perform a qY2H affinity ladder analysis. Some sub-ensembles (*e.g.*, **5**) lead to a correct order of the PPIs according to their affinities, but several sub-ensembles gave wrong results (**1**, **8**, **10**). *B*. When the ten sub-ensembles were combined to a single ensemble of one million cells, a correct affinity ladder was obtained. *C*. The affinity ladders obtained from three subsequent ensembles of one million cells are presented. They all show the same correct order as in Fig. 4.
